# Prodiginine Production in *Streptomyces coelicolor* Correlates Temporally and Spatially to Programmed Cell Death

**DOI:** 10.1101/240689

**Authors:** Elodie Tenconi, Matthew F. Traxler, Charline Hoebreck, Gilles P. Van Wezel, Sébastien Rigali

**Affiliations:** InBioS - Centre for Protein Engineering, Institut de chimie B6a, University of Liège, B-4000 Liège, Belgium; Department of Plant and Microbial Biology, Berkeley University of California, CA 94720, United States.; Molecular Biotechnology, Institute of Biology, Leiden University, PO Box 9505, 2300RA, Leiden, The Netherlands

**Keywords:** cell death and differentiation, antitumor antibiotics, bacterial development, Confocal laser microscopy

## Abstract

Programmed cell death (PCD) is a common feature of multicellularity and morphogenesis in bacteria. While cell death has been well documented when *Streptomyces* species switch from vegetative (nutrition) to aerial (reproduction) growth, lethal determinants are yet to be discovered to unveil the genetic basis of PCD in mycelial bacteria. In this work we used prodiginines of *Streptomyces coelicolor* as model to test the hypothesis that a bacterium uses ‘self-made’ antiproliferative DNA-damaging agents as toxins of their PCD process. Spatio-temporal visualisation of the autofluorescence of prodiginines reveals that their biosynthesis is triggered in the dying zone of the colony prior to morphological differentiation of the mycelium. A prodiginine nonproducer showed hyper-accumulation of viable filaments, with increased RNA and proteins synthesis when most of the mycelium of the wild-type strain was dead when prodiginine accumulated. Addition of a prodiginine synthesis inhibitor also strongly favoured viable over dead filaments. As self-toxicity has also been reported for other producers of DNA-damaging agents we propose that cytotoxic metabolites synthetized during the morphological transition of filamentous bacteria may be used to execute PCD.

**Significance Statement:** Actinobacteria are prolific producers of compounds with antiproliferative activity, but why these bacteria synthetize metabolites with this bioactivity has so far remained a mystery. Using prodiginines (PdGs) as model system, we revealed that the spatio-temporal synthesis of these molecules correlates to cell death of the producer *Streptomyces coelicolor* and that inhibition of their synthesis results in hyper-accumulation of viable filaments. Since PdGs potentiate death of *S. coelicolor* recurrently prior to morphological differentiation, this is a form of programmed cell death (PCD). Hence, next to weapons in competition between organisms or signals in inter- and intra-species communications, we propose a third role for secondary metabolites i.e., elements required for self-toxicity in PCD processes.

## Introduction

Traditionally, bacteria were regarded as unicellular microorganisms that rapidly grow and divide via binary fission. The concept of multicellularity among prokaryotes was recognized only three decades ago, and is particularly evident in actinobacteria, cyanobacteria and myxobacteria (1, 2). These are bacteria with a complex life cycle and morphological and chemical differentiations that are switched on in response to environmental signals. A hallmark of multicellular organisms is programmed cell death or PCD (1, 3, 4). PCD is the cell cycle-dependent autolytic dismantling or *cannibalism* of older cells, and serves to produce the building blocks required for a new round of macromolecule synthesis to sustain developmental growth, such as mycelia or fruiting bodies. In *Bacillus. subtilis*, sporulation is preceded by a form of PCD known as cannibalism in a subpopulation of the biofilm (5, 6). In *Myxococcus*, three subpopulations that show division of labour arise, with cells differentiating into spores or into peripheral rods, while the remaining cells undergo PCD (7). In the cyanobacterial *Anabaena*, cell death is controlled by the circadian rhythm, which implies careful programming (8).

Several rounds of PCD occur as part of the complex developmental program of the mycelial *Streptomyces* (9–13). Streptomycetes are filamentous Gram-positive high G+C bacteria that reproduce via sporulation. Their life cycle starts with a spore that germinates to grow out and form a mycelium consisting of multinucleoid vegetative hyphae. When reproduction is required, the older mycelium is used as a substrate to form aerial hyphae, which differentiate into chains of unigenomic spores. This morphological differentiation is accompanied by chemical differentiation, whereby many natural products, including antibiotics, anticancer compounds and antifungals, are produced. The lytic degradation of the vegetative mycelium is a clear manifestation of PCD, presumably to provide the nutrients necessary to produce the reproductive aerial hyphae (10, 11, 14, 15).

The molecular triggers and genetic basis for PCD in mycelial bacteria are still largely unknown. ‘Antibiotics’ should be regarded as serious candidates. Antibiotic resistance is indeed mandatory for antibiotic makers such as streptomycetes, which infers that their ‘weapons’ against competitors are also active against the producing species (16–21). To survive production of their natural products, *Streptomyces* possess self-defense mechanisms identical to those developed and acquired by human and animal pathogens (22–24). Self-toxicity - and therefore self-resistance - extends well beyond antibacterial agents; for example, anticancer molecules damaging DNA are also deadly to the producer (25). Mechanisms of resistance to antitumor antibiotics that destroy DNA have been identified in almost all producers and include i) drug sequestration (25–27), ii) drug inactivation (28–30), iii) drug efflux (29, 31–34), and iv) target repair or protection (35–38).

Amongst the *Streptomyces* molecules currently explored in human disease therapy, the tripyrrole red-colored prodiginines or prodigiosin-like pigments (PdG) have gained interest due to their promising antitumor, immunosuppressive and anti-inflammatory properties (39–42). Their topoisomerase-inhibiting activity by intercalating DNA explains why prodiginines are toxic to so many different organisms and also display antimalarial, anthelmintic, antifungal, and antibacterial activities (43). Such a broad spectrum of activity suggests that the PdG-producing species should be sensitive to their own molecules. Indeed, in *Streptomyces coelicolor* the onset of expression of the *red* cluster (encoding the PdG biosynthetic pathway) coincides with the entrance of this species into a period of growth cessation – the so-called transition phase (44–50). In addition, prodiginine production coincides with a round of massive cell death in the course of which the *Streptomyces* multicellular filamentous network undergoes drastic morphological changes associated with the sporulation process. Time-space monitoring of *redD* expression, the activator of PdGs *red* biosynthetic genes, is also confined to ageing, lysed filaments (48). Finally, prodigiosin-like pigments are induced by stressing culture conditions like excess of metal ions and pH shock (51, 52), co-cultivation with competing microorganisms (53, 54), and feeding the medium with dead bacterial cells (55).

In the light of the known high cytotoxicity of prodiginines, it is remarkable that the DNA-damaging PdGs are not secreted by *S. coelicolor*, but rather accumulate internally, right at the time of growth cessation. This seemingly suicidal and at the same time well-programmed mechanism suggests that PdGs may play a role in the control and/or progress of PCD. An advantage of using PdGs as our model to correlate anticancer production to PCD is that the timing and localization of their production is easily monitored *in situ* due to their red autofluorescence (56). In this work we demonstrate that prodiginine production correlates to dying filaments in time and space and that absence of PdGs reduces cell death in *S. coelicolor*, resulting in hyper-accumulation of viable filaments. We propose that prodiginines and, in general, intracellular DNA-damaging metabolites are main protagonists of PCD processes in the producing organisms.

## Results

### Prodiginine biosynthesis is concomitant in time and space with PCD

In order to define if prodiginines (PdGs) are associated with PCD in *S. coelicolor* we evaluated if their production correlates in time and space with the part of the culture that undergoes the massive round of vegetative mycelium degradation that precedes the onset of the reproductive aerial growth. PdG production was monitored throughout the life cycle of *S. coelicolor*, making use of their red autofluorescence (RAF) (56). Spores (10^7^ cfu) of *S. coelicolor* M145 were spread onto the surface of R2YE agar plates, and 0.5-mm thick slices of confluent solid cultures were collected at different time points and imaged via confocal fluorescence microscopy. *In situ* visualization of PdG production (under non-saturated excitation conditions) revealed that weak RAF appeared at the surface of the vegetative mycelium at around 36 h, and that this signal reached its maximum level at 50 h (Figure 1a, Figure S1). From that time point onwards, the RAF intensity decreased abruptly to persist at approximately one third of its maximal intensity (Figure S1). The accumulation of PdGs was therefore maximal (~50 h) at the morphological transition phase, before the vegetative mycelium differentiates into aerial hyphae (between 50 and 64 h, Figure S1).

**Figure 1.**
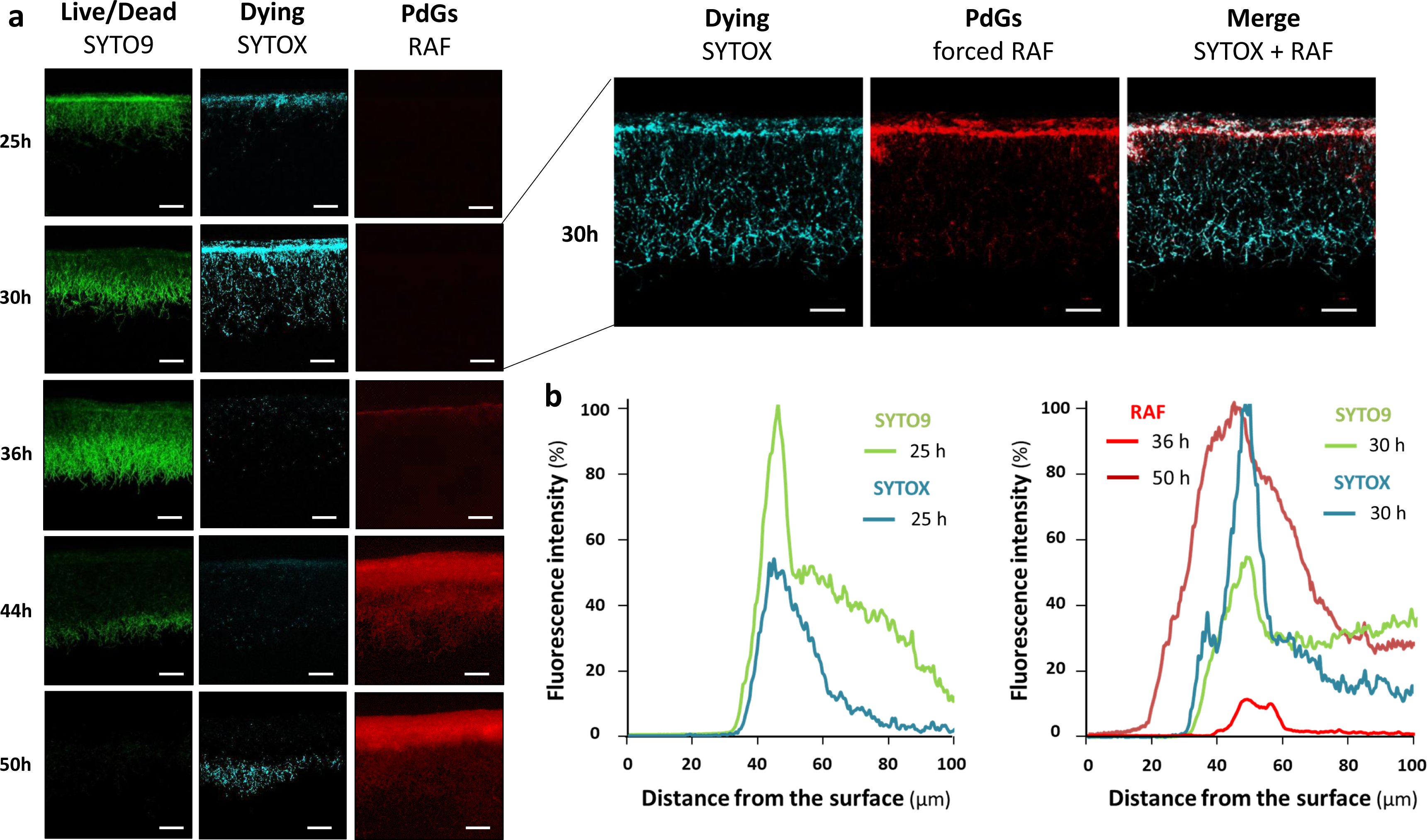
Spatio-temporal occurrence of production of PdGs and cell death in *S. coelicolor*. (**a**) Images represent cross-sections of confluent lawns of *S. coelicolor* M145 grown on R2YE agar plates. Maximal fluorescence intensity associated with PdG biosynthesis (RAF), dying filaments (SYTOX-staining, blue-colored), or live/dead filaments (SYTO9-staining, green colored), was reached after 50, 30, and 25 h of growth, respectively. The voltage on the photomultiplier tubes (PMT) for RAF was fixed to 600 V. On the right panels the PMT was increased to 680 V in a 30 h old cross section in order to visualize RAF showing early PdG production that starts from the thin band of cell death. Bars, 40 µm. (**b**) Quantification of RAF, SYTO9, and SYTOX fluorescence signals across the first 100 µm of the *S. coelicolor* culture. Note that the peak of maximum RAF, SYTO9, and SYTOX occurred in the same thin band at a distance of about 50 µm from the surface of the culture.

Simultaneous *in situ* quantification of RAF and dying cells revealed that the membrane-damaged filaments, which are permeable to SYTOX, reached their highest level at around 30 h of culture, about 20 h before the maximal intensity of RAF (~50 h) (Figure 1a and 1b). This maximum of accumulation of dying filaments occurred ~5 h after the peak of SYTO9 fluorescence (~25 h), which, at time points prior to PdG production (see below), stains both live and membrane damaged (dying) cells (Figure 1a and 1b). Quantification of SYTO9 and SYTOX signals along the first 100 µm of the *S*. *coelicolor* culture revealed that both dyes present a peak of fluorescence at the same distance of ~50 µm from the surface of the culture (Figure 1b). This rigorous spatial co-localization at an approximately 5 h interval suggests that dying filaments observed at 30 h emanated from those presenting the highest SYTO9 signal at 25 h. Quantification of PdGs along the same vertical axis showed that the earliest RAF signal detected at 36 h (under non-saturated excitation conditions Figure 1a and 1b) also arises at the same distance of ~50 µm from the surface of the culture, right after the maximum of accumulation of dying cells (Figure 1b). Increasing the voltage on the photomultiplier tubes (PMT) during imaging of 30 h old samples revealed that the forced early RAF signal matched the thin 10 µm band of the culture where first dying filaments were seen (Figure 1a, right panel) supporting the idea that PdG biosynthesis spatially coincides with cell death but would be posterior to it.

At a later time point (40 h) qualitative analysis of PdG fluorescence and accumulation of dying cells (SYTOX stained) along the vertical axis of a confluent solid culture showed a clear correlation between RAF and the zone of cell death visible in the upper part of the culture (Figure 2a). Conversely, no or extremely weak RAF was seen in SYTO9 stained sections, and *vice versa* (Figure 2a). The fact that maximum RAF (50 h) coincided with i) the lowest amount of SYTO9 (live and dead) staining, and ii) maximal SYTOX (dead) staining (Figure 3) further supports that PdGs might be associated with the massive round of cell death observed at the surface of the culture and preceding the morphological differentiation of *S. coelicolor*. To further demonstrate that PdGs accumulate in the dead zone at the surface of the *S. coelicolor* culture, we repeated the monitoring of dying cells using propidium iodide (PI) as alternative fluorescent dye for staining membrane-damaged filaments. PI displays maxima of excitation/emission of red fluorescence very close to the maxima of RAF of PdGs (56). Monitoring PI fluorescence will thus also reveal RAF when PdGs are produced. At 30 h of growth, prior to PdG production, PI display fluorescence at the upper part of the culture (Figure 2b) with a pattern similar to the one observed with SYTOX staining (Figure 1a). At later time points, when PdGs are produced, the fluorescence of PI in the upper part of the culture (Figure2b) is observed in the same zone as where RAF associated with PdGs was observed (Figure 1a), the fluorescence signal being increased when both RAF and PI were recorded. That RAF recorded together with PI fluorescence staining dying filaments display similar patterns as those recorded for RAF alone further supports that PdGs accumulate in the dead zone at the surface of the culture. However, it has to be noted that, as observed for SYTOX, PI also stains damaged filaments deeper inside the agar plate where RAF from PdGs is not observed (Figure 2).

**Figure 2.**
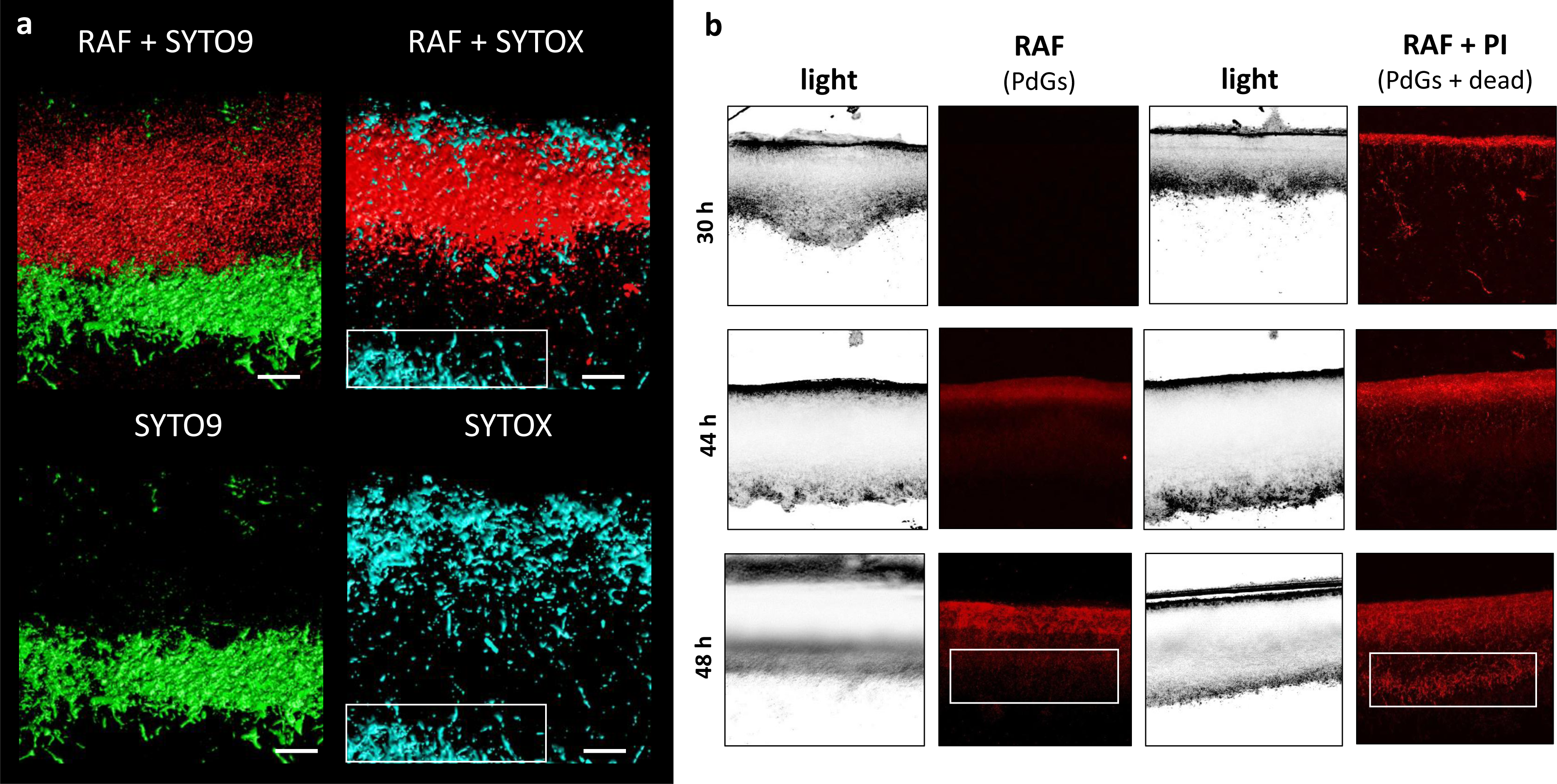
Localization of the production of prodiginines in the dying zones of the *S. coelicolor* culture. (**a**) Fluorescence confocal micrographs (3D reconstruction) of two transverse sections of an *S. coelicolor* M145 culture grown for 40 h on R2YE agar plates. Left panel: RAF (red-colored) and staining of viable filaments (green-colored, SYTO9). Right panel: RAF (red-colored) and staining of dying filaments (blue-colored, SYTOX). Bar, 20 µM. (**b**) Visualization of the red fluorescence associated with PdG production (RAF) and propidium iodide staining (PI). White rectangles indicate the zone where SYTOX (a) and PI (b) stained dying filaments do not match with PdG production.

### PdGs have anti-proliferative activity

Since PdG production apparently correlated to the zone of the culture where cell death occurred (Figure 2), we attempted to monitor their production at the filament size scale. Repetitive assays revealed that, at time points where *S. coelicolor* abundantly produces PdGs (from 40 h onwards), SYTO9 and SYTOX hardly stained any of the cells that display RAF. Visualisation of filaments that display both RAF, and SYTOX or SYTO9 fluorescence was only possible at lower mycelial density (below the zone of high fluorescence signals in the first 50-60 µm), or at the brief moment in the life cycle (30 h) when the level of PdGs is still low inside the filaments (Figure 3). Specimens where we could observe intracellular PdG production inside filaments stained with SYTO9 or SYTOX are presented at Figure 3. However, once PdG accumulated above a certain level inside the filaments, SYTOX and SYTO9 failed to stain, leading to partial or complete ‘ghost’ filaments that could only be visualised by RAF (Figure 3). This inhibition of SYTO9 and SYTOX staining is most likely the result of the competing DNA-intercalation by PdGs and/or because of the PdGs-induced destruction of nucleic acids.

**Figure 3.**
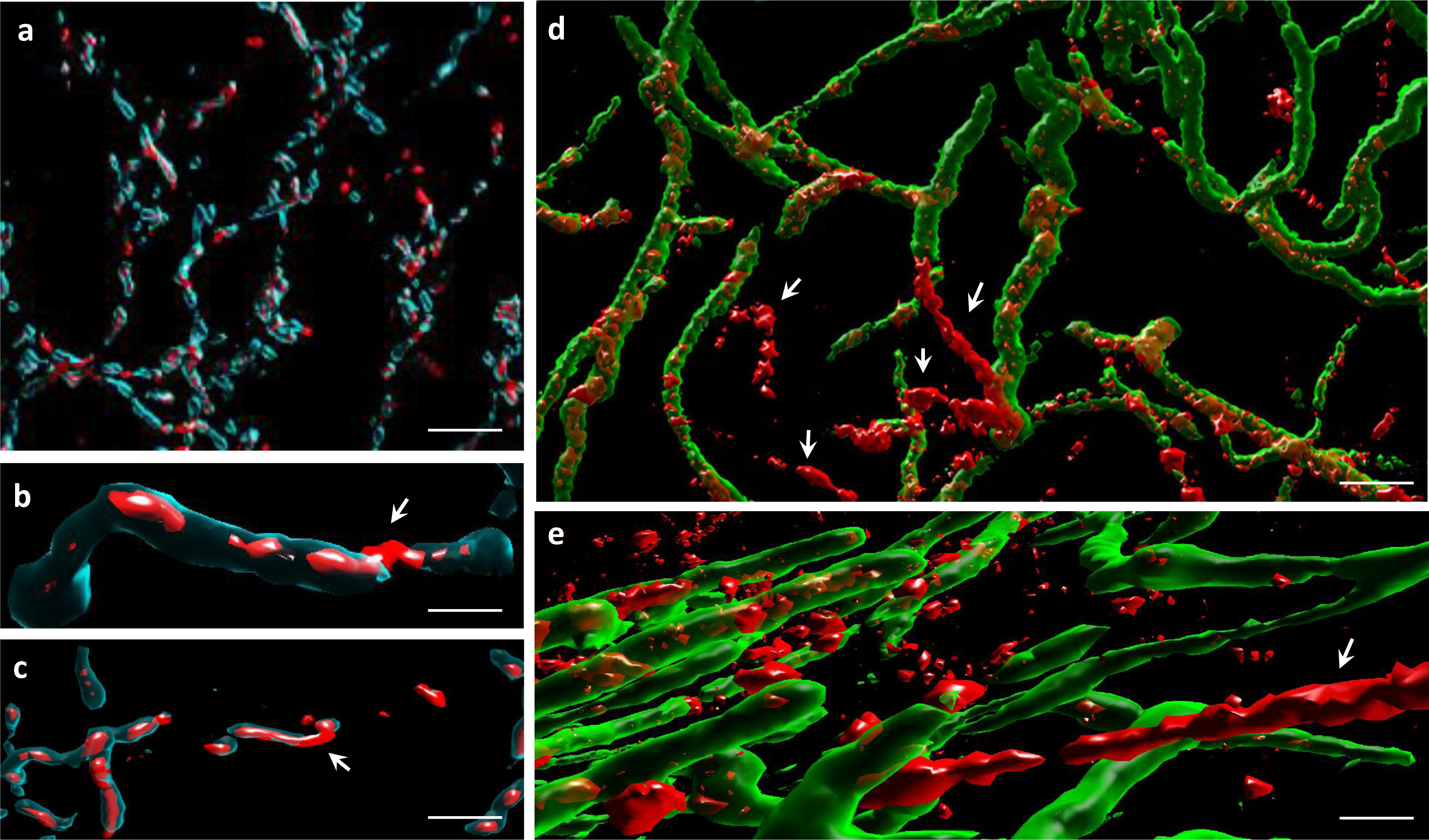
Monitoring of PdG production at the filament size scale in *S. coelicolor*. (**a** to **c**) Filaments of *S. coelicolor* M145 inoculated on R2YE plates displaying both RAF and SYTOX (blue) staining. White arrows point to portion of filaments not stained with SYTOX and with abundant PdG production visualized by RAF. Bars, 10 µm (a), 2 µm (b), and 5 µm (c). (**d** and **e**) Filaments of *S. coelicolor*M145 inoculated on R2YE plates displaying both RAF and SYTO9 (green) staining. White arrows point to ‘ghost filaments’ with abundant production of PdGs and therefore not stained with SYTO9. Bars, 2 µm (e), and 1 µm (f).

The observed lack of staining with SYTOX or SYTO9 once the RAF signal is high, and therefore when PdG production abundantly accumulated, led us to hypothesize that these DNA-damaging metabolites that remain intracellular, may be involved in the destruction of the DNA in the vegetative mycelium of *S. coelicolor*. To investigate this, we first compared the accumulation of viable and dying filaments between *S. coelicolor* M145 and its *redD* null mutant M510, which fails to produce PdGs (Figure 4). The *redD* null mutant also displayed a band of dying hyphae at the upper surface of the culture at 30 h (Figure 4), which may be seen as an argument that the production of PdGs is a consequence, and not the cause, of the onset of cell death. However, at 50 h of growth, which corresponds to the time point of maximal RAF intensity in the wild-type strain (Figure 1 and S1), the PdG-nonproducing strain M510 still displayed many SYTO9-stained cells, while the parental strain M145 showed only very weak staining along the vertical axis of the culture (Figure 4). Importantly, we did not observe any fluorescence when SYTOX was added after 50 h of growth in the *redD* null mutant (Figure 4), suggesting that all filaments stained by SYTO9 were viable. The absence of SYTO9 and SYTOX staining in the zone of the culture that displayed maximal RAF suggests that PdGs had caused widespread DNA destruction within the vegetative mycelium. Complementation of the *redD* mutant restored PdG production and resulted in the subsequent massive loss of filaments stained with SYTO9 (Figure S2).

**Figure 4.**
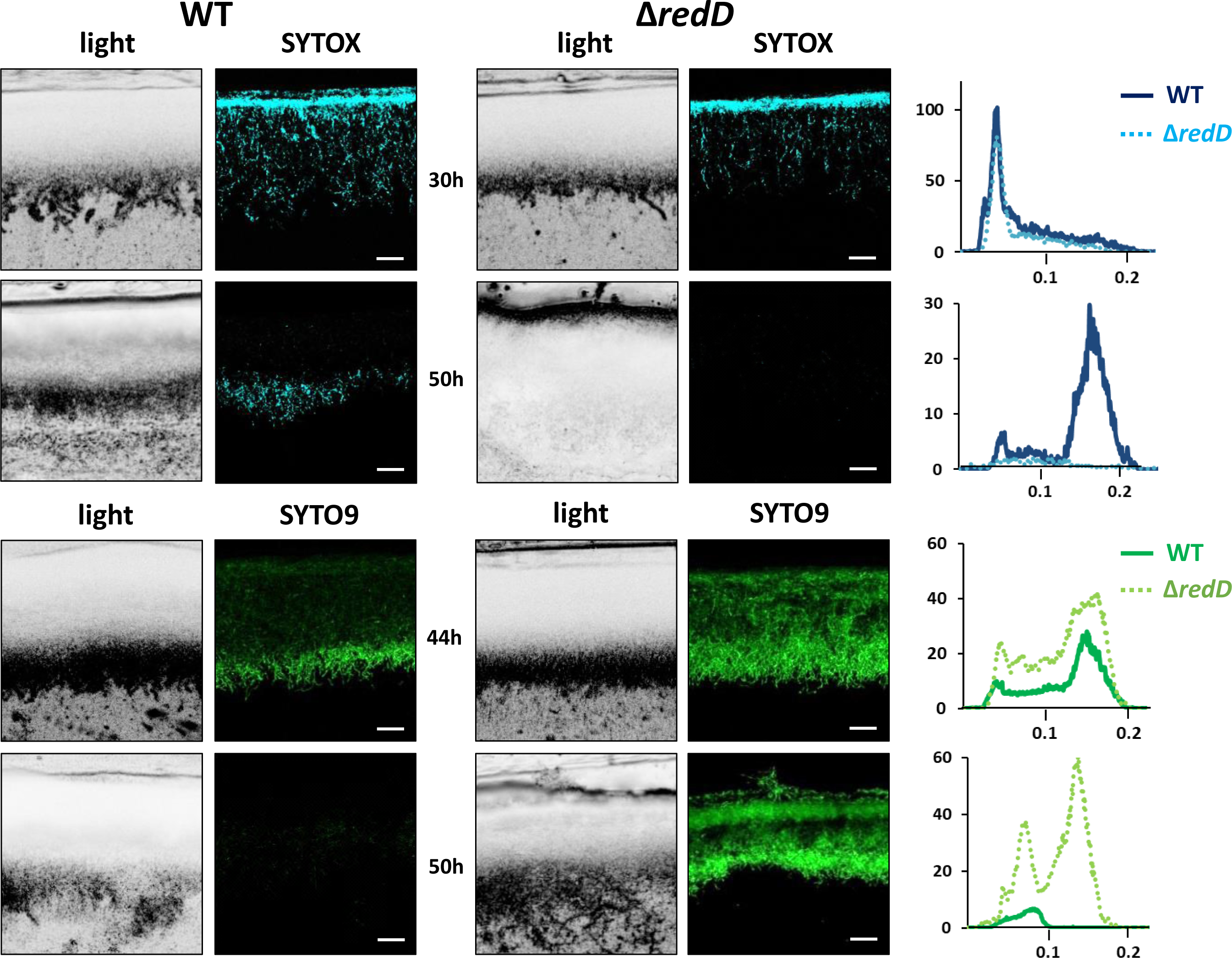
Prodiginine biosynthesis causes massive loss of viable filaments. SYTOX (blue) and SYTO9 (green) staining of cross sections of culture of *S. coelicolor* M145 (wild-type, WT) and its *redD* null mutant inoculated in R2YE agar plates. Bars, 40 µm. Note that (i) the PCD round prior to PdG production also occurs at 30 h in the *redD* null mutant of *S. coelicolor*, and (ii) the higher accumulation of viable filaments (only stained by SYTO9) in the *redD* mutant. Quantification of the SYTOX and SYTO9 fluorescence signals are displayed in plots next to the microscopy pictures.

Additionally, we monitored and compared other markers of viability and metabolic activity between the wild-type strain M145 and the *redD* mutant, namely RNA and protein synthesis (Figure 5a and 5b). For this purpose, R2YE agar plates were covered with cellophane discs prior to inoculation with spores of either *S. coelicolor* M145 or its *redD* mutant M510, to allow collection of the biomass from the spent agar. The presence of the membranes on the top of the plates caused accelerated production of PdGs, with a peak in RAF intensity 10 h earlier than observed in plates not covered with cellophane discs (40 h instead of 50 h) (Figure S3a and S1). Visualization of total intracellular RNA synthesis showed a drop in the accumulation of 16S and 23S rRNA in the wild-type strain right after the peak of PdG biosynthesis (Figure S3b). In contrast, after that time point rRNA levels remained significantly higher in the *redD* mutant as compared to the parental strain (Figure S3b). Assessment of total protein content of the intracellular crude extracts also revealed distinct profiles between the two strains, with the *redD* mutant displaying higher amounts throughout the time course as compared to the parental strain, which showed a drastic drop of protein accumulation just after the peak of RAF at 40 h (Figure S3c). The higher intracellular accumulation of macromolecules (DNA, RNAs, and proteins) strongly suggests that the metabolism of the PdG-nonproducer remains active at stages of the life cycle where the large majority of the vegetative mycelium of the wild-type strain is dead.

**Figure 5.**
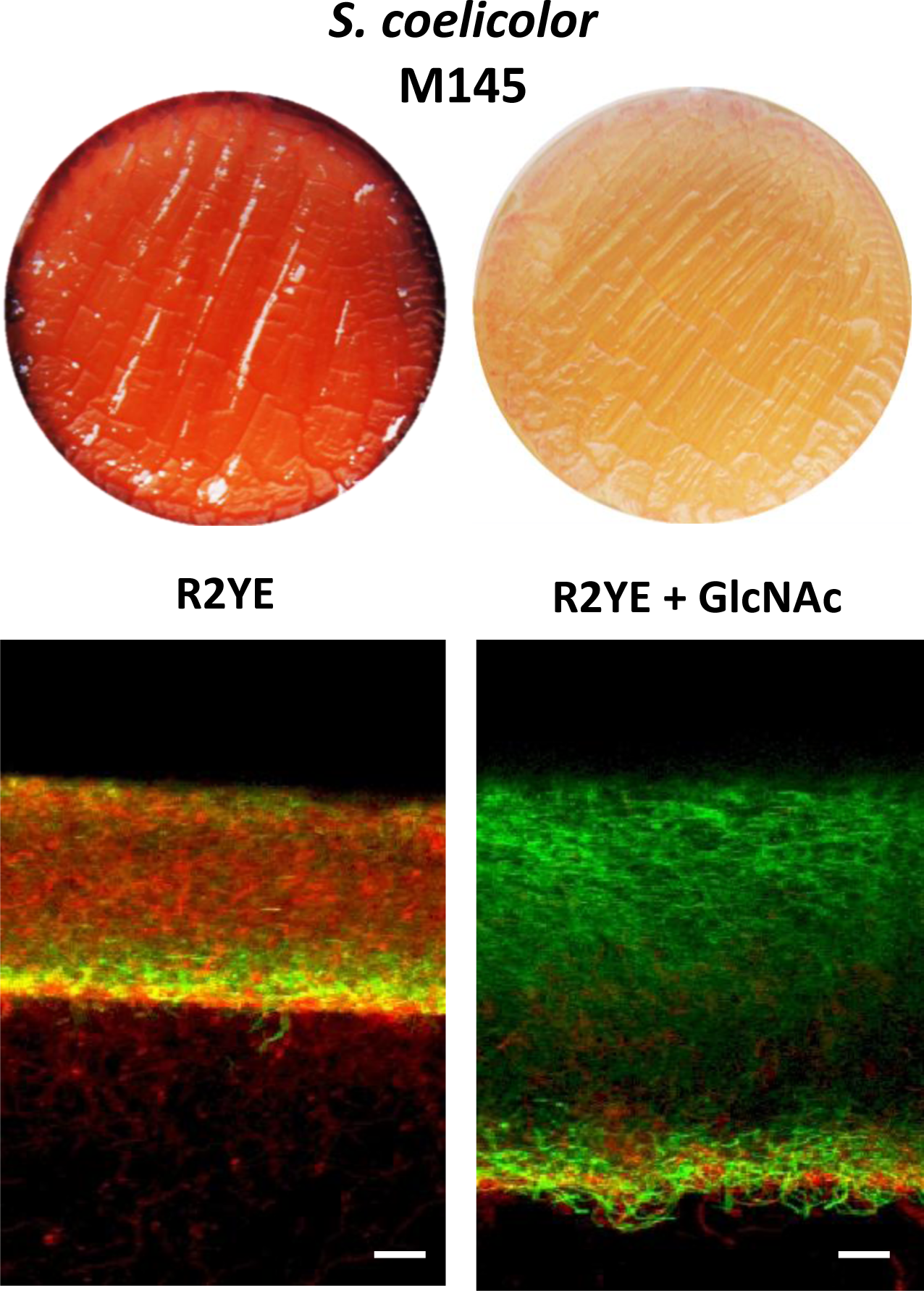
Addition of the PdG biosynthesis inhibitor GlcNAc results in accumulation of viable filaments. Effect of GlcNAc (1%) on SYTO9 (green) filament staining, and PdG and PI (red) fluorescence. Bars, 20 µm. *S. coelicolor* M145 was inoculated in R2YE agar plates.

### Inhibiting conditions for PdG production reduces cell death

To further test the idea that *S. coelicolor* itself is vulnerable to intracellular PdGs and that PdG are induced by cell death, we used a culture condition known to reduce or inhibit PdG production in *S. coelicolor* and assessed if it also correlates with reduced cell death and/or higher accumulation of viable filaments. As a first assay, 2.10^7^ spores of *S. coelicolor* M145 were used to inoculate the R2YE solid medium with or without addition of the aminosugar N-acetylglucosamine (GlcNAc) known to block morphogenesis at the vegetative state and PdG production in *S. coelicolor* grown on rich media (57, 58) (Figure 5). *In situ* visualization of samples of confluent solid cultures collected at 42 h revealed that the presence of the morphogenesis-blocking agent GlcNAc resulted in a massive accumulation of SYTO9 stained filaments (Figure 5a). As GlcNAc containing samples were devoid of the red fluorescence associated with PdG production and PI staining (Figure 5), it suggests that all SYTO9 stained filaments were viable. In contrast, without GlcNAc, the mycelium of *S. coelicolor* displayed important red fluorescence (from both PI and PdG) associated with dying cells and presented a much lower growth as deduced from the thickness of the mycelial culture (Figure 5). This suggests that inhibiting or reducing PdG synthesis correlates with higher accumulation of viable filaments.

## Discussion

Do natural products with anti-proliferative activities, such as PdGs, participate in the PCD process of the producing microorganism? This is the motivating question of the work we present here, which aimed to assess if DNA-damaging compounds like PdGs, which are promising drugs in cancer therapy, also display anti-proliferative activities in the producing organism, in this case *S. coelicolor*. As PdGs are not secreted but only accumulate internally in *S. coelicolor*, this hypothesis is directly in line with the concept of these molecules being auto-destructive rather than acting on competing microbes. We found that the production of PdGs perfectly coincides in time and space with a cell death event that kills the overwhelming majority of the *S. coelicolor* colony. This broad and rapid destruction of the mycelium is assisted by the PdGs as in the PdG non-producing strain (the *redD* null mutant) cell death was attenuated, and RNA and protein synthesis continued at time points where normally these macromolecules are destroyed in the parental strain. Loss of cell death and accumulation of viable filaments were also observed when the culture medium was supplemented with the PdG production inhibitor GlcNAc.

### Cell death is the trigger of PdG biosynthesis

Next to phenotypic characterization of mutants, a complementary means to uncover the role(s) played by a secondary metabolite in nature is to identify signals that regulate its biosynthesis (i.e. ‘*what controls you will tell us who you really are*’). When we visualized the cultures at an early time point, we found that the biosynthesis of PdGs is not triggered homogeneously but is instead spatially restricted to a very thin (<10 µM) band in the upper part of the *S. coelicolor* culture (Figure 1). This thin band of PdG production arises exactly where a round of cell death is constantly observed at the same moment of the life cycle of *S. coelicolor*. This observation corroborates previous findings that *redD* expression occurred only when a proportion of the hyphae had lysed, suggesting that transcription of *redD* is confined to ageing vegetative mycelium (48). It has also been demonstrated that the production of PdGs of *S. coelicolor* is induced by the addition of dead cells in the culture medium (55). The initial band of cell death where the production of PdGs started also corresponds to the thin band where we observed maximum SYTO9 and SYTOX staining at 25 h and 30 h, respectively (Figure 1b). PdG biosynthesis thus arises where the density of the population is maximal and therefore could be governed by a quorum sensing regulatory pathway. That synthesis of killing factors (PdGs) follows in time with the zone of the culture with the highest density of the population is most likely not just a coincidence. The high density observed at 25 h in the thin band in the upper part of the culture suggests highly active metabolism during aerobic growth with important production of reactive oxygen species (ROS) which could be the cause of the massive cell death observed in the same zone at 30 h (Figure 1) and proposed as triggered for the formation of the aerial mycelium of *Streptomyces* (59).

### What would be the evolutionary advantage of producing cytotoxic compounds prior to morphogenesis?

For *Streptomyces*, it is generally accepted that the death of the vegetative mycelium provides nutrients for the advancement of later stages of the life cycle, *i*.*e*., the erection of aerial hyphae and their subsequent maturation into spore chains (11, 12, 14, 15). However, we note that there is a current paucity of data that actually support this claim. At the same time, the toxicity of PdGs might also prevent scavenging of these newly liberated nutrients by other bacteria and/or fungi. The model above (*i*.*e*. chemical protection of its own food reservoir) is appealing as it rationalizes why a PCD event might precede morphogenic development and why a small molecule that is also toxic to other organisms might be an ideal mediator of this process. In this case, the killing agent is a structurally complex compound whose synthesis requires a large cluster of 23 genes. Making PdGs might seem an expensive strategy to participate in a cell death process. But making a complex natural product is only ‘expensive’ if the evolutionary pay-off for making it is modest. However, if the pay-off is large, then it is a favourable strategy. So, for the PdGs if the pay-off is ensuring its own food reservoir during the sporulation process, then it might not be expensive at all.

### Specific and controlled suicide

A microorganism that triggers its own cell death in a controlled manner has to make sure that (i) not all cells are killed by the process and (ii) it has to sense that the killing molecule is ‘home made’. In other words, the compartments close to the future site of development have to undergo cell death in a controlled manner. It is important to note that the production of PdGs is indeed well controlled, and strictly correlates to the so-called transition phase in submerged cultures, and to the phase immediately preceding the onset of morphological development in solid-grown cultures (44, 46, 48). This strongly suggests that the cell death caused by the PdGs is indeed programmed, in other words represents PCD in the true sense, as a way to eradicate the old mycelial biomass for the next growth phase, namely the aerial hyphae and spores. One could easily imagine that the physiological reaction to sensing the presence of a toxic compound must be very different if the molecule originates from competitors that share the same environmental niche or, instead, if the molecule is self-made. This also explains why microorganisms that undergo PCD to sustain metamorphosis would use their own molecule for signalling the timing of differentiation. If they would all use the same trigger their development would be synchronized, while their fitness to the environmental is different. Similarly, plants all have their own cocktail of hormones and physico-chemical parameters for inducing flowering. In addition, having a killing system not ‘universal’ (conserved molecule or mechanisms in all actinobacteria) allows also having its own mechanism of resistance (see examples of specific resistance to enediyene, daunorubicin and doxorubicin, mitomycins and bleomycins in the introduction). Therefore, a secret code in the form of a complex natural product with resistance is required.

However, a key experimental result presented here suggests that the reality is more complicated; the *redD* null mutant (which fails to produce PdGs) still undergoes an initial, limited round of PCD. Importantly, there are also many culture conditions where *S. coelicolor* does not produce PdGs but still undergoes PCD, which means that cell death can be completed in the absence of PdGs. When PdG biosynthesis is required remains unclear, but seems inextricably linked to growth conditions where microorganisms are facing challenging conditions. Indeed, previous works both in *Streptomyces* and *Serratia* species, reported that PdG biosynthesis is associated with diverse, stress-inducing culture conditions such as UV-light exposure, oxidative stress, nutrient depletion or competition with other microorganisms (43, 50, 53, 55). In many cases authors have postulated that the ecophysiological role of PdGs is to protect the producing strain against these various stresses (43). How do the cell-eradicating properties of PdGs described in this work, and those also well documented for eukaryotic cell death or apoptosis (42), fit with the proposed protective roles in various stressful conditions? Again, when part of the colony is challenged by lethal stress conditions, inducing controlled cell death in that part of the colony would be a good strategy to spare at least part of the population. In such a scenario, at least a fraction of the population might survive for dissemination to more appropriate living conditions.

### Concluding remarks and perspectives

The data presented here shows that PdGs are closely associated with programmed cell death process of the producing organism. Specifically, an initial phase of PCD appears to serve as a ‘detonator’ that triggers the widespread self-killing of the *S. coelicolor* biomass through the action of the PdGs. Our work highlights for the first time genes encoding toxic determinants of the PCD phenomenon in mycelial bacteria. Providing answers to how some filaments survive the production of these lethal compounds and switch to a reproductive life-style are key questions going forward. Finally, the proposed role in PCD of antitumor antibiotics could be played by other types of antibiotics (targeting the cell wall, the translational machinery,…) in species that do not produce DNA-damaging metabolites. Hence, next to weapons in warfare between organisms or molecules involved in inter- and intra-species communications, our work proposes a third role for secondary metabolites that is elements required for self-toxicity in PCD events either necessary to maintain an appropriate growth balance, to cope with stress conditions, or as part of the developmental programme.

## Methods

### Strains and culture conditions

*S. coelicolor* M145 and its *redD* mutant M510 (60) were used as the wild-type strain and as PdGs non-producer, respectively. R2YE (61) agar plates were used for phenotypic characterization. Where applicable, R2YE plates were covered with cellophane membranes (GE Osmonics Labstore, Ref K01CP09030). Plates were inoculated with 500 µl of a 2.10^7^ cfu/ml spore suspension.

### Complementation of the *redD* mutant M510

The *S. coelicolor redD* mutant M510 was complemented by introducing plJ2587 harbouring *redD* (SCO5877) with its native upstream region. A DNA fragment containing the *redD* upstream region (567 bp) was generated by PCR using primers (5’- GAATTCCCCCTGCTGCTCCAGGG −3’) and (5’- GGATCCCCCAATATGTTGATTTCCACGC −3’) with engineered EcoRI and BamHI sites, respectively, and cloned into pJET1.2 (Thermo Fisher Scientific). After sequence confirmation, the fragment was retrieved through EcoRI and BamHI restriction digest, gel purified, and cloned into an EcoRI/BamHI-linearized plJ2587 (62) upstream of *redD* resulting in plasmid pELT003. The complementation construct, as well as the empty plJ2587 plasmid, were introduced into the *redD* mutant through intergeneric conjugation as described previously (63). All thiostreptone resistant colonies transformed with pELT003 (gene *tsr* in plJ2587) presented the intracellular red pigmentation and red fluorescence associated with PdG production confirming the complementation of the *redD* mutant phenotype.

### *In situ* visualization by confocal fluorescence microscopy

Samples were prepared as described previously (56). Samples were examined under a Leica TCS-SP2 and Leica TCS-SP5 confocal laser-scanning microscopes. SYTO9 and SYTOX stained samples were examined at a wavelength of 488 for excitation and 530 nm (green) for emission. Red autofluorescence of PdGs and propidium iodide-stained samples were examined at a wavelength of 543 nm (Leica TCS-SP2) or 568 nm (Leica TCS-SP5) for excitation and 630 nm (red) for emission as described previously (64). Image processing and 3D reconstruction of *Streptomyces* filaments were performed as described previously (56). Z-Stacks of confocal images (objective HCX PL APO 63 × 1.20 W CORR UV and pixels 512 × 512) were processed using the Fiji (ImageJ) software. For analyses of dense culture across a transversal section, ten Z-stack images from a 238.1 μm section were used for a standard deviation Z-projection (for light images) or a maximal Z-projection after applying a Gaussian Blur filter with a radius of 1 (for RAF images). For the 3D confocal image stack reconstitution, UCSF Chimera software was used to visualize 84 z-stack images from a 85.2 μm section (objective HCX PL APO 63 × 1.20 W CORR UV with a zoom of 2.8 and pixels 512 × 512) preprocessed by the Fiji software (Gaussian Blur filter for RAF image stack).

### RNA and proteins extraction and quantification

Total intracellular RNA was extracted using the phenol/chloroform/isoamyl alcohol protocol, mainly as described previously (65). The *S. coelicolor* mycelium was scraped with a spatula from the R2YE agar plates covered with cellophane discs, put into 2 ml tubes and frozen at −70 °C. 50 mg of mycelium were first subjected to lysozyme digestion (2 mg/ml final concentration in 2 ml of extracted buffer pH8), and then incubated with proteinase K (0.5 mg/ml; 1h at 55 °C) prior to nucleic acids extraction. DNA was removed from total nucleic acids extracted by using the kit Turbo DNA-free (Ambion). Proteins were extracted from 50 mg of mycelium as described previously (56) after sonication using 30 second pulse for 10 minutes (Bioruptor, Diagenode, Liège, Belgium) in 500 µl of extraction buffer. Protein concentration was determined by measuring absorbance at 280nm.

## Acknowledgements

ET work was supported by a FRIA grant and by the Belgian program of Interuniversity Attraction Poles initiated by the Federal Office for Scientific Technical and Cultural Affairs (PAI no. P7/44). SR is Maître de Recherche at FRS-FNRS. We are thankful to Prof. Patrick Motte from InBioS - PhytoSystems and Centre for Assistance in Technology of Microscopy (CATµ) of the ULiège, and to Dr. Sandra Ormenese and Jean-Jacques Goval from the Cell Imaging and Flow Cytometry GIGA Platform at ULiège. This work is dedicated to the memory of Martine Bansse-Tenconi (1952-2013).

---

**Figure S1.**
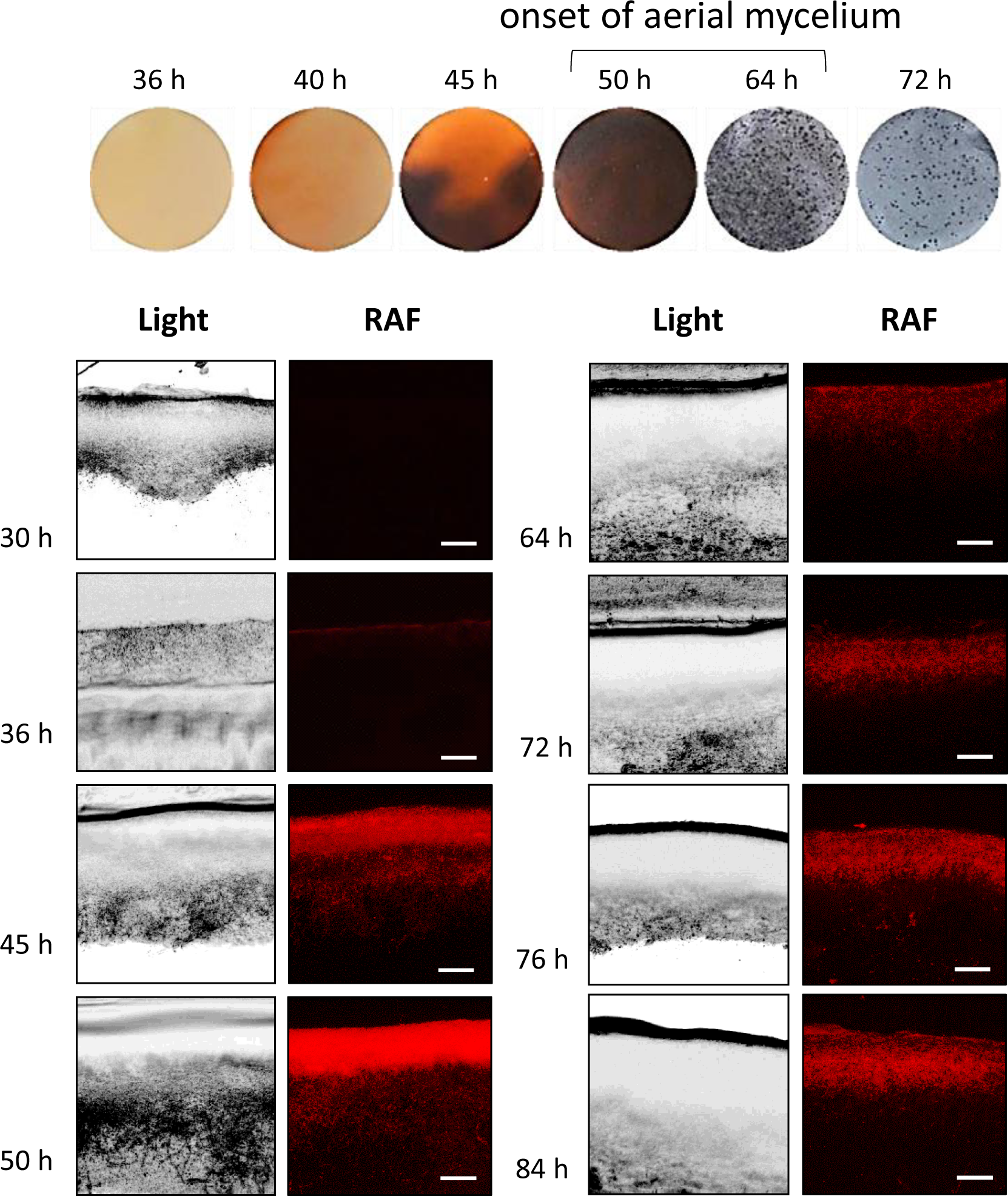
Visualization and quantification of RAF throughout the life cycle of *S. coelicolor*. *in situ* visualization of RAF by confocal microscopy in cross section of a confluent *S. coelicolor* culture on R2YE agar plates. Bars, 40 µm. Phenotypes of *S. coelicolor* grown in R2YE agar plates are shown to visualize the delay between the onset of PdG production and the onset of the morphological differentiation (white mycelium appearing on the surface of the plate).

**Figure S2.**
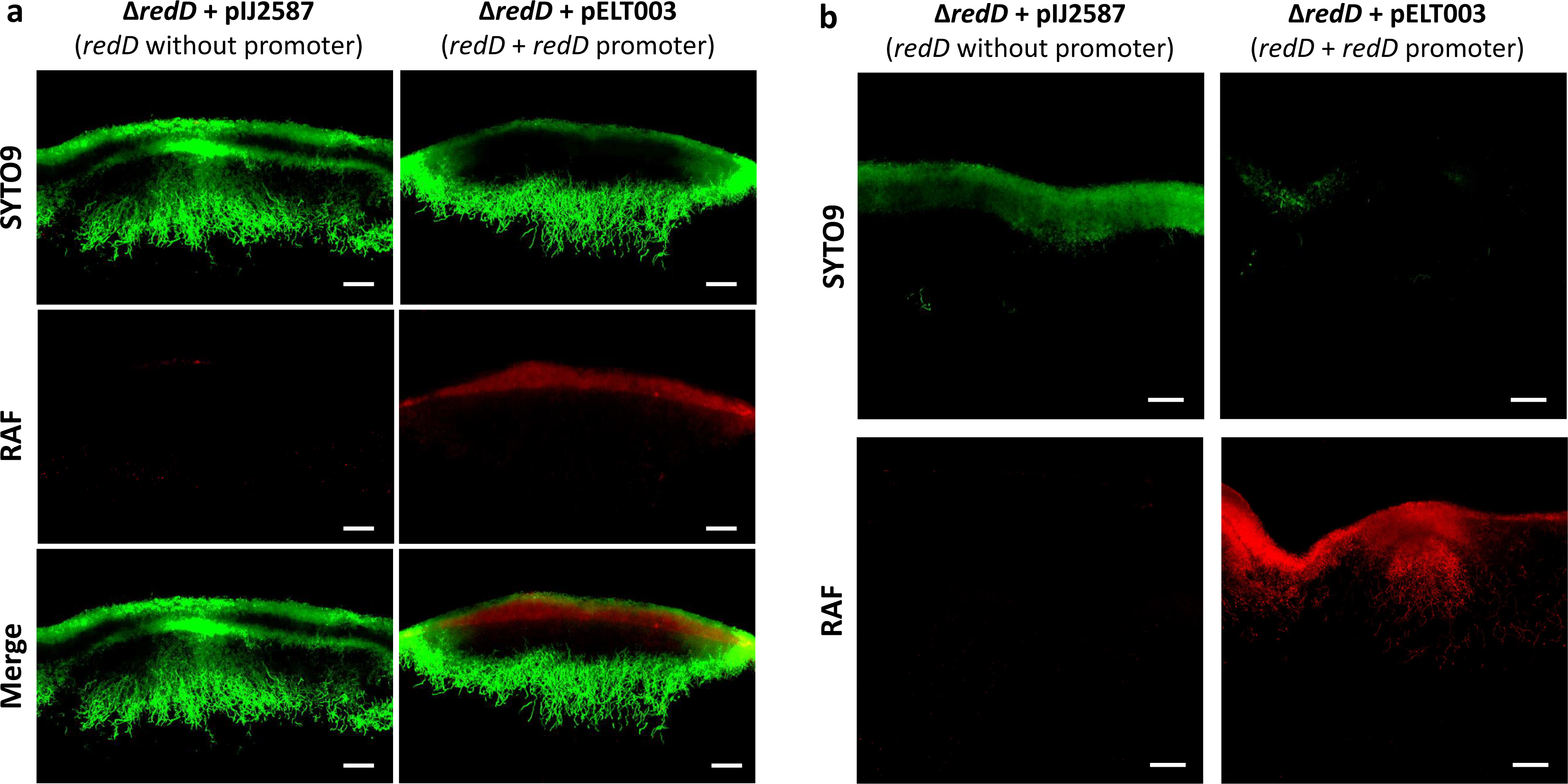
Complementation of the *redD* null mutant restores PdG production and loss of SYTO9-stained viable filaments. (a) Effect of the complementation of the *redD* null mutant at the colony size scale. Bars, 20 µm. (b) Effect of the complementation of the *redD* null mutant on cross sections of 50 h old cultures. Bars, 40 µm.The *redD* null mutant was complemented with pELT003 containing an entire copy of *redD* with its own promoter (567 bp upstream of the *redD* start codon). Plasmid plJ2587 (62) containing *redD* without its upstream region (no promoter) was used as control.

**Figure S3.**
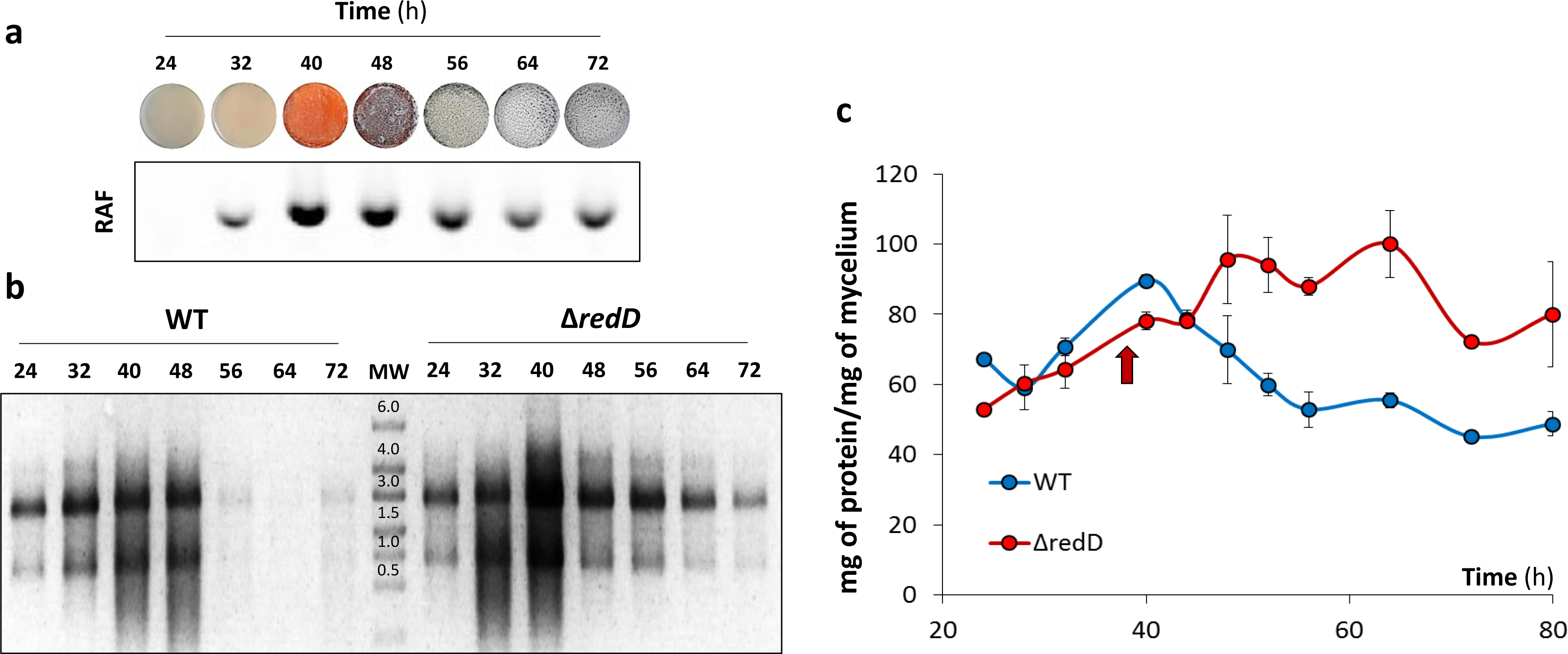
Effect of the deletion of *redD* on RNA and protein synthesis. (**a**) PdG production throughout the life cycle of*S. coelicolor*. RAF from mycelium extracts of *S. coelicolor* grown on the R2YE medium covered with cellophane discs. Extracts were deposited in an agarose gel and RAF was monitored after migration as described previously (56). (**b**) Ethidium bromide stained agarose gels showing intracellular RNA extracted from *S. coelicolor* M145 (WT) and its *redD* null mutant (∆*redD*). (**c**) Quantification of the total protein content in crude extracts of mycelia of *S. coelicolor* WT (M145) and its *redD* null mutant. The red arrow indicates the timing of maximal PdGs biosynthesis.

